# Gene pathogenicity prediction of Mendelian diseases via the Random Forest algorithm

**DOI:** 10.1101/553362

**Authors:** Sijie He, Weiwei Chen, Hankui Liu, Shengting Li, Dongzhu Lei, Xiao Dang, Yulan Chen, Xiuqing Zhang, Jianguo Zhang

**Affiliations:** BGI Education Center, University of Chinese Academy of Sciences, Shenzhen 518083, China; BGI-Shenzhen, Shenzhen 518083, China; Shenzhen Key Laboratory of Neurogenomics, BGI-Shenzhen, Shenzhen 518083, China; China National GeneBank, BGI-Shenzhen, Shenzhen 518120, China; Center of Prenatal Diagnosis, ChenZhou No.1 People’s Hospital, Hunan 423000, China

**Author notes:** These authors contributed equally to the work (Sijie He, Weiwei Chen). Corresponding authors: Jianguo Zhang, Xiuqing Zhang. **Conflict of Interest:** The authors declare that they have no conflict of interest.

**Keywords:** Machine learning, Mendelian disease genes, Gene pathogenicity prediction, Gene dominance prediction

## Abstract

The study of Mendelian diseases and the identification of their causative genes are of great significance in the field of genetics. The evaluation of the pathogenicity of genes and the total number of Mendelian disease genes are both important questions worth studying. However, very few studies have addressed these issues to date, so we attempt to answer them in this study.

We calculated gene pathogenicity prediction (GPP) score by a machine learning approach (random forest algorithm) to evaluate the pathogenicity of genes. When we applied the GPP score to the testing gene set, we obtained accuracy of 80%, recall of 93% and area under the curve (AUC) of 0.87. Our results estimated that a total of 10,399 protein-coding genes were Mendelian disease genes. Furthermore, we found the GPP score was positively correlated with the severity of disease.

Our results indicate that GPP score may provide a robust and reliable guideline to predict the pathogenicity of protein-coding genes. To our knowledge, this is the first trial to estimate the total number of Mendelian disease genes.

## Introduction

In recent years, the identification of pathogenic genes for Mendelian diseases (MD) has developed rapidly. Wenger *et al.* indicated that an average of 266 OMIM phenotypes with known molecular bases and 241 gene-disease associations for MD are reported annually (Wenger et al. 2017). Currently, more than 5000 single-gene disorders have been reported, and nearly 4000 genes are responsible for them. The key points in MD research are how to efficiently evaluate the pathogenicity of each gene and how many genes among the ~20,000 protein-coding genes may cause MD.

Variant-level prediction methods are widely used in the identification of MD genes. There are dozens of tools for performing variant-level prediction, such as SIFT (Ng et al. 2003), Polyphen (Adzhubei et al. 2010), GERP++ (Davydov et al. 2010), MutationTaster (Schwarz et al. 2014), CADD (Kircher et al. 2014), REVEL (Ioannidis et al. 2016), and so on. In contrast to the numerous variant prediction tools, only a few tools focus on gene-level prediction, such as RVIS (Petrovski et al. 2013), DNE (Samocha et al. 2014), EvoTol (Rackham et al. 2015) and GDI (Itan et al. 2015), which may play an irreplaceable supplementary role in MD gene identification. These existing gene-level prediction tools rely mainly on a single characteristic of genes, which is the intolerance to functional variants, to make predictions. Although the genic intolerance values produced by these tools may reflect the pathogenicity of genes, they didn’t give a cutoff for deeming them pathogenic.

The pathogenicity of genes is decided by more than one kind of factor. If we combine multiple characteristics instead of just one characteristic to do the prediction, we may have a better result. Machine learning is an efficient method for classification and prediction. It may combine many features from the studied objects and provide a comprehensive judgment of new objects. The development of variant-level prediction tools indicates the ones using machine learning tend to show better performance. The random forest algorithm is a well-developed model of machine learning that can handle multiple features and tolerate samples with missing features. This algorithm is very suitable for predicting the pathogenicity of genes.

In this study, we applied a machine learning approach (random forest algorithm) that combined 201 gene-level characteristics to produce the gene pathogenicity prediction (GPP) score. The GPP score showed better performance than residual variation intolerance score (RVIS) and gene damage index (GDI) in distinguishing MD genes. Our results estimated that a total of 10,399 protein-coding genes were MD genes. The characteristics of GPP score were also analyzed. Gene dominance prediction (GDP) score and gene recessiveness prediction (GRP) score were calculated by the same method to evaluate the inheritance model of MD genes. The 3 scores were integrated in a list called the gene catalog of Mendelian diseases (GCMD). Our results may be applied to MD research, especially for the identification of pathogenic genes. This is the first trial to provide a clear cutoff for judging MD genes and to estimate the total number of them.

## Methods

### Data collection and gene standardization

The data contained several gene sets extracted from ClinVar (Landrum et al. 2018), OMIM (http://omim.org/), MalaCards (Rappaport et al. 2017), the “super hero project” (Chen et al. 2016) and other sources; variant data extracted from 1000 Genomes Project (KG) (Sudmant et al. 2015) and Genome Aggregation Database (containing whole-genome data (GAD) and whole-exome data (ExAC)) (Lek et al. 2016); gene-level characteristics extracted from Refgene, STRING (Szklarczyk et al. 2017), HomoloGene, GTEx, ANNOVAR (Wang et al. 2010); and other sources. The annotation results of variants and genes were involved as well. Data from different sources may have inconsistent gene names, which may lead to some confusion. In our study, all the gene names from different data sets were standardized according to the HUGO Gene Nomenclature Committee (HGNC) (see Supplementary methods).

### Gene set selection and gene-level characteristic filtration in machine learning approach

To explore a comprehensive method to predict the pathogenicity of genes, we applied a machine learning approach. The gene sets and gene-level characteristics used in machine learning were produced as follows.

Gene set selection: The loss-of-function (LOF) variant tolerant genes were obtained from the KG, GAD and ExAC databases, among which 630 high-quality genes were used as the training set of negative genes, and the remaining 850 genes were used as the testing set of negative genes. To ensure a balance between the number of positive and negative genes used in our model, we extracted and selected 630 and 850 MD genes as the training and testing sets of positive genes, respectively.

Gene-level characteristic filtration: We extracted many gene-level characteristics from ANNOVAR, STRING, Refgene, HomoloGene, GTEx and DOMINO (Quinodoz et al. 2017), as well as several characteristics calculated by ourselves, including the gene intolerance scores, gene potential damaging scores, and so on. In total, we enrolled 405 characteristics. If some genes were missing a value for a characteristic, we used the median value of other genes to fill it. We excluded genes with missing values for more than 50% of all characteristics. Then, we performed some filtration and finally kept 201 characteristics (Supplementary **Table 1**).

The details are provided in the Supplementary methods.

### Gene pathogenicity prediction score produced by random forest algorithm

After the gene sets and gene-level characteristics were obtained, we adjusted the parameters of the random forest algorithm (ntree and mtry) and built a model with the training gene sets and gene-level characteristics. We applied the model to the testing gene set to evaluate its effect. The 10 most important characteristics of the model were analyzed. Then, the GPP score of all protein-coding genes were calculated by the model, the number of predicted pathogenic and non-pathogenic genes (Np and Nn) of MD were obtained according to their scores (the default cutoff 0.5 was used). Considering the false-positive rate (FPR) and false-negative rate (FNR), we estimated the number of MD genes (Nm) as: *Nm = Np* ×(1 − *FPR*)+ *Nn × FNR*. We then collected several disease gene sets and analyzed the prediction accuracy and score distribution of them. The detailed gene sets and the gene selection method are listed in Supplementary **Table 2**.

### Prediction of dominant and recessive models of Mendelian disease genes

To distinguish dominant genes and recessive genes, we applied the same pipeline we used when calculating the GPP score to calculate the GDP and GRP scores, respectively. Briefly, we collected genes from the autosome that showed dominant, recessive and both inheritance patterns. In our model, 1243 positive genes and 1584 negative genes were used for the GDP score calculation, while 1985 positive genes and 842 negative genes were used for the GRP score calculation. After the filtration, 183 gene-level characteristics were kept. We did not divide the genes into a training and a testing set, and we applied 4X cross-testing to do the calculation (see Supplementary methods).

The 2 scores (GDP and GRP) and GPP score are integrated in Supplementary **Table 3**.

## Results

### The performance of GPP score produced by the random forest algorithm

We applied a machine learning approach (random forest algorithm) to calculate gene pathogenicity prediction (GPP) score. After the adjustment of parameters, 201 gene-level characteristics were used, and we found ntree=500 and mtry=46 to be suitable parameters. When we applied the GPP score produced by this model to the testing gene set, we obtained accuracy of 80%, recall of 93%, FPR of 26% and FNR of 10% (the default cutoff 0.5 was used). The receiver operating characteristic (ROC) curve showed an area under the curve (AUC) of 0.87. We compared the performance of the GPP score with that of RVIS and GDI on the testing gene set and found that the GPP score performed significantly better than the other two tools (**Fig 1A**). The 10 most important characteristics identified by the mean decrease in the Gini coefficient of the model mainly belonged to 3 categories (gene intolerance scores, variant damage prediction results and gene interaction score) (**Fig 1B**). Then, we evaluated the performance of each characteristic on the testing gene set. The ROC curve showed that the AUC ranged from 0.5 to 0.76 (Fig 1C), which was much lower than that of the GPP score.

**Fig 1.**
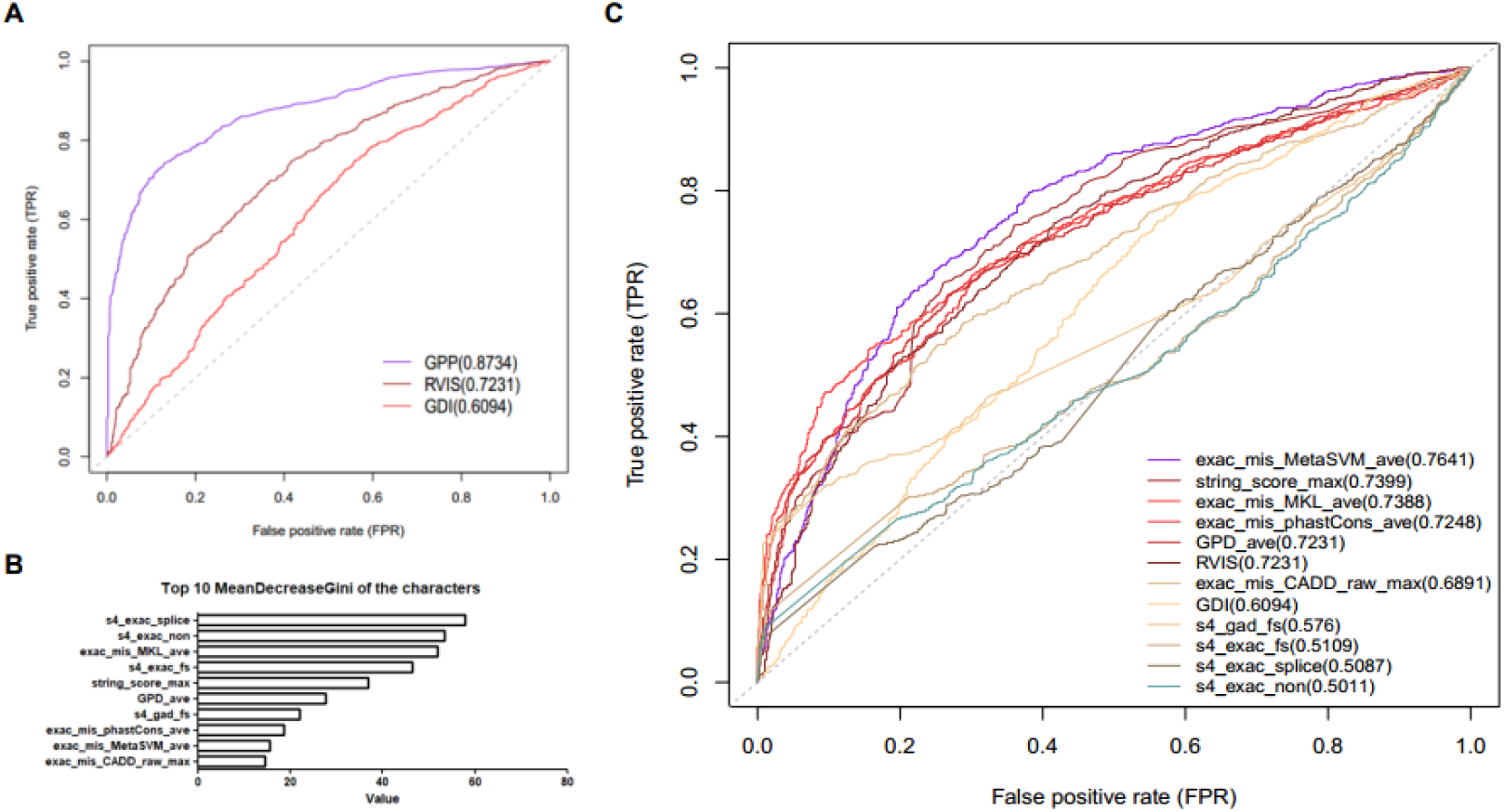
The ROC curve of GPP score and the 10 most important characteristics of the model. A. The ROC curve on the testing gene set. The AUCs of the GPP score and two similar tools, RVIS and GDI, are 0.87, 0.72 and 0.61, respectively. B. The 10 most important characteristics identified by the mean decrease in the Gini coefficient, ranked by their values. C. The ROC curve on the testing gene set. The AUCs of the 10 most important characteristics are listed on the right, ranked by their values.

We collected reported MD genes and susceptible genes from OMIM, LOF variants intolerant genes from ExAC and known pathogenic genes from several resources to evaluate the prediction accuracy of the GPP score. The genes without GPP score were excluded. The cutoff to distinguish pathogenic and non-pathogenic genes of MD was set as 0.5. MD genes showed higher accuracy (92.8%) than that of susceptible genes (74.5%), LOF variants intolerant genes showed even higher accuracy (96.7%), and pathogenic genes showed a bit lower accuracy (87.4%) (Table 1). It was comprehensible that susceptible genes showed lower prediction accuracy because part of them were not MD genes. The accuracy of pathogenic gene set was not that high because a part of pathogenic genes of non-Mendelian diseases were involved. And the results indicated that LOF variants intolerant genes in ExAC may contain massive potential MD genes waiting to be confirmed.

**Table 1.**
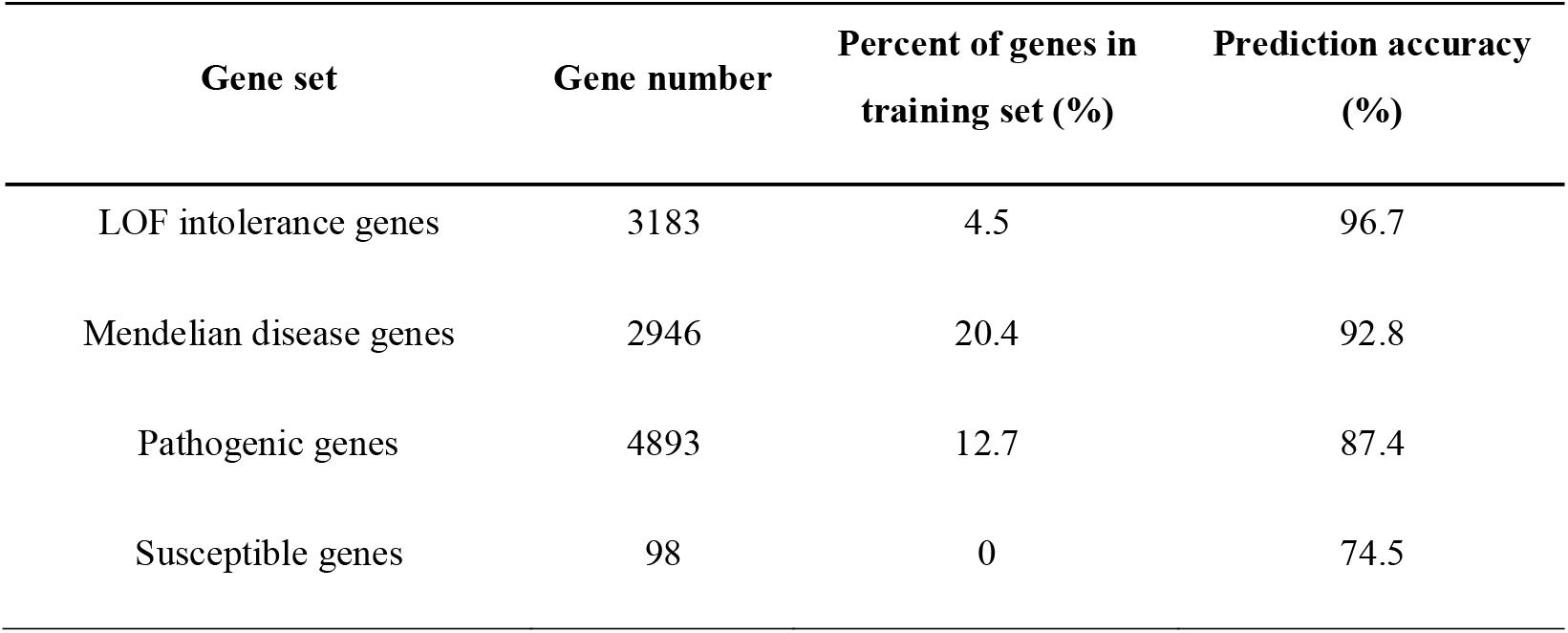
Prediction accuracy of GPP score for several kinds of pathogenic gene set

### The estimation for the number of Mendelian disease genes and the GO analysis

Among 18,226 high-quality protein-coding genes involved in our model, 13,401 genes were predicted to be pathogenic (the default cutoff 0.5 was used). Considering the FPR and FNR of our model, we estimated that approximately 10,399 genes were MD genes.

We extracted pathogenic and non-pathogenic genes of MD according to our prediction result and reported MD genes from OMIM, then we performed gene ontology (GO) analysis of them. Totally 2984 reported pathogenic genes of Mendelian diseases (reported_PM), 13401 predicted pathogenic genes of Mendelian diseases (predicted_PM) and 4825 predicted non-pathogenic genes of Mendelian diseases (predicted_NM) were involved. The number of genes with at least one GO entry of these gene sets were 2974, 13154 and 4377, respectively. We calculated the average number of GO entries per gene and the values of the three gene sets were 22.9, 17.2 and 8.5, respectively.

In general, the reported_PM and the predicted_PM sets showed little difference, while the predicted_NM set showed larger difference compared to them (Supplementary Fig 1). We found that the predicted_PM and predicted_NM sets showed little difference compared to the reported_PM set in some GO pathways, such as membrane part (GO:0044425), extra-cellular region (GO:0005576), immune system process (GO:0002376), and so on. In particular, the predicted_NM set showed higher percentage of enrichment compared to the other two sets in two pathways, which were signal transducer activity (GO:0004871) and cell killing (GO:0001906).

### GPP score is positively correlated with the severity of disease

Two peaks were observed in the GPP score (**Fig 2A**). The proportion of overlap between the genes with different GPP scores and reported MD genes was analyzed. We found that genes with higher scores showed a higher proportion of overlap with reported MD genes, and the proportion ranged from 4.4% (GPP scores under 0.5) to 30.8% (GPP scores above 0.9) (**Fig 2B**). We also examined the distribution of predicted and reported MD genes on each chromosome, but we did not find obvious hot or cold spots (Supplementary **Fig 2**).

**Fig 2.**
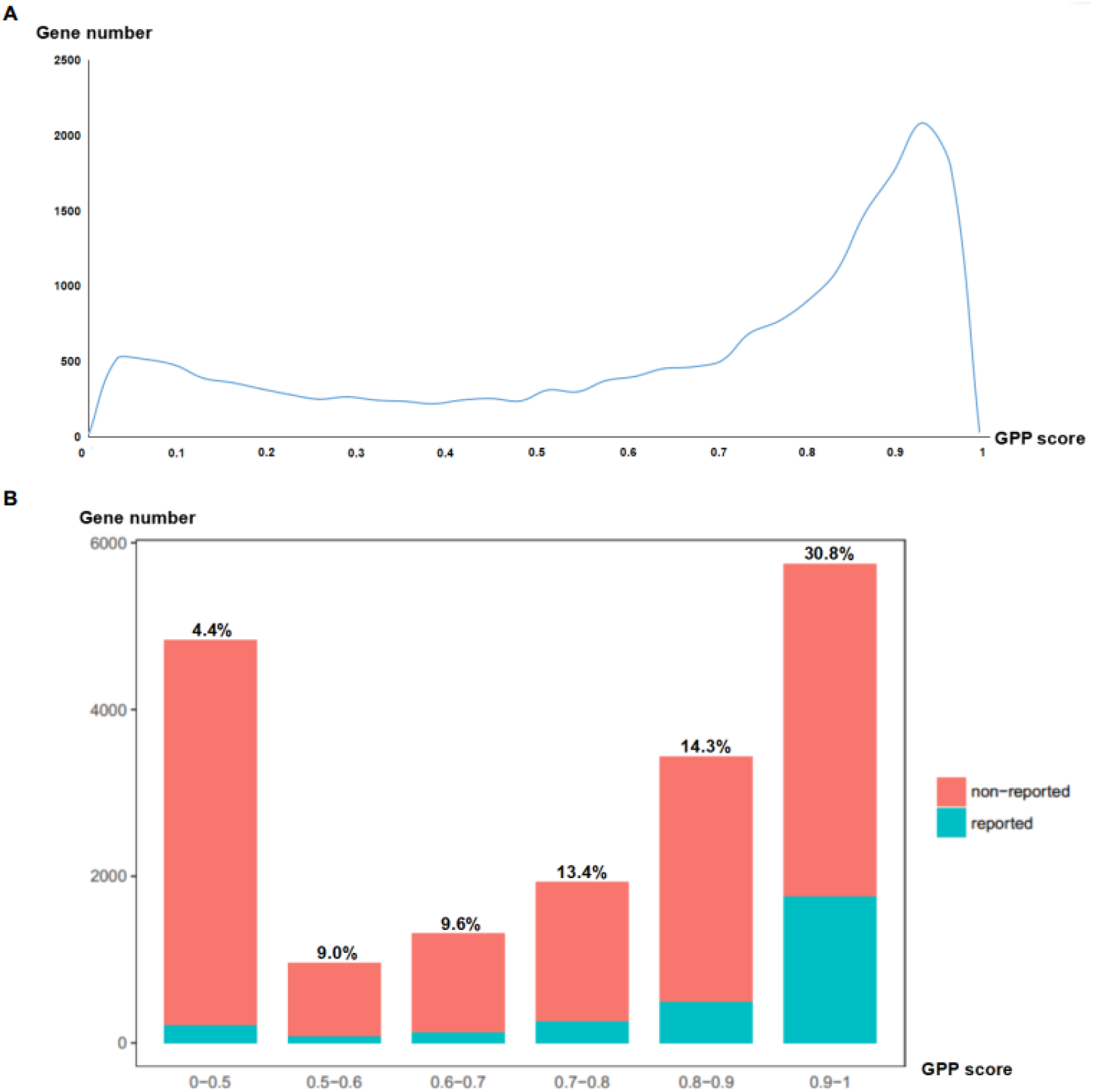
Distribution of the GPP score and proportion of reported Mendelian disease genes for different scores. A. The distribution of the GPP score of all protein-coding genes shows 2 peaks, and the majority of genes are predicted to be MD genes. The X-axis represents the GPP score, while the Y-axis represents the number of genes. B. Genes are divided into 6 sets by the GPP score. Each bar consists of two parts. The blue part shows reported MD genes, and the red part shows non-reported ones. The proportion of reported genes is listed at the top of each bar.

The distribution of the GPP score for several kinds of pathogenic gene set was observed. We found that genes with known inheritance models showed significantly higher scores than susceptibility genes (**Fig 3A**). And we found the scores of LOF genes were higher than those of GOF genes (Wilcoxon rank sum test, p≤0.001) (**Fig 3B**). To determine whether the GPP score could reflect the age of onset or severity of the disease, we examined the score distribution of several pathogenic gene sets accordingly. Among several kinds of neurological disease, intellectual disability (ID) and autism spectrum disorder (ASD) tended to show an early age of onset, the onset of schizophrenia (SCZ) mainly occurred in late adolescence, Alzheimer’s disease (AD) often occurred in the elderly, and epilepsy (EP) occurred in a wide range of age. The scores of these gene sets showed little difference (**Fig 3C**). The pathogenic genes of diseases with severe phenotypes (neuromuscular disease, metabolic disease, congenital heart disease and hereditary tumors) showed higher scores, while the genes of diseases with mild phenotypes (skin diseases such as psoriasis and ichthyosis) showed lower scores. These results indicated GPP score was positively correlated with the severity of disease. Genes of complex diseases (hypertension, obesity and diabetes) and male infertility showed scores between those of severe and mild diseases (**Fig 3D**).

**Fig 3.**
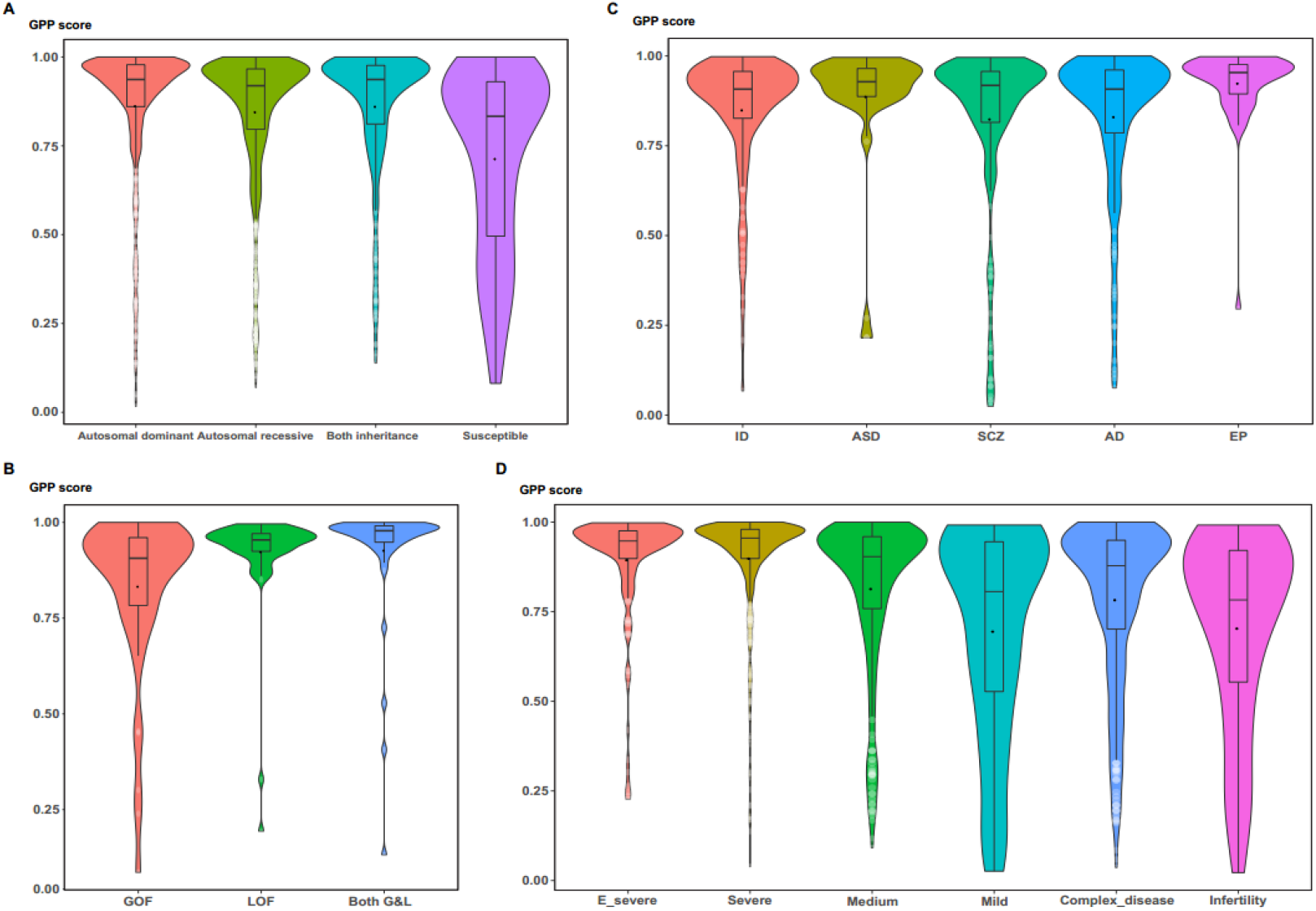
Distribution of the GPP score for different pathogenic gene sets. A. The distribution of genes with different inheritance models, including dominant, recessive, both dominant and recessive genes in autosome and susceptible genes. B. The distribution of GOF, LOF and both G&L genes. C. The distribution of pathogenic genes of neurological diseases with different ages of onset: intellectual disability (ID), autism spectrum disorder (ASD), schizophrenia (SCZ), Alzheimer’s disease (AD) and epilepsy (EP), from left to right. D. The distribution of genes responsible for diseases with different severities. E_severe set contains genes of some early-onset severe Mendelian diseases, severe set contains genes of diseases with severe phenotype (neuromuscular disease, metabolic disease, congenital heart disease and hereditary tumors), medium set contains genes of some diseases with medium phenotype (ophthalmic diseases and hearing disorders), mild set contains genes of some diseases with mild phenotype (skin diseases such as psoriasis and ichthyosis), complex_disease set contains genes of some complex diseases (hypertension, obesity and diabetes) and infertility set contains genes of male infertility.

### The dominant and recessive patterns for pathogenic genes

We applied the same machine learning approach to calculate GDP score and GRP score to predict dominant and recessive genes. After adjustment of the parameters, we obtained accuracy of 75%, recall of 64%, FPR of 24%, FNR of 25% and AUC of 0.82 for the GDP score and accuracy of 81%, recall of 97%, FPR of 20%, FNR of 15% and AUC of 0.808 for the GRP score. We also checked the score distribution of all MD genes and found that more genes tended to follow the recessive model (Supplementary **Fig 3**). Our prediction results showed that there were 4942 autosomal dominant genes and 10,041 autosomal recessive genes. Considering the FPR and FNR, we estimated that the numbers of real dominant and recessive genes were 5871 and 8537, respectively.

## Discussion

Our results estimate that 10,399 genes are MD genes, which is much more than currently reported number. This is the first time to estimate the total number of MD genes. The estimated number may indicate that there are many pathogenic genes (or lethal genes) of MD waiting to be discovered. Lethal genes are those with important function and the dysfunction of them will cause death before birth. Lethal genes are rarely identified in disease research, so we wanted to provide a cutoff of the GPP score for lethal genes. We extracted some lethal genes from the Mouse Genome Informatics (MGI) database. The quartile value of them was 0.888, which may suggest a cutoff of lethal genes.

Analysis of the score distributions of different gene sets showed that the GPP score was positively correlated with disease severity. Infertility is a special kind of disease, which may have a strong influence on the next generation but little influence on the patients themselves. We also analyzed the diseases with different ages of onset and found no significant difference between the scores of these gene sets. These results indicate that the GPP score may reflect disease severity, where severity means the degree of threat to the health or survival of the individuals themselves but not the next generation. The gene sets we used may contain some non-Mendelian disease genes that influence the results. So, we exclude genes not in the reported MD gene pool and do the analysis again, and similar results are obtained.

In this study, we ultimately calculated 3 gene-level scores to evaluate the pathogenicity of all protein-coding genes and to assign the dominant or recessive model to each MD gene. The GPP score may help to identify MD genes, while the GDP and GRP scores may help to identify which genes follow dominant or recessive models. To explore a wider range of application of our scores, we applied the GPP score to some genes related to schizophrenia identified by GWAS (Li et al. 2017), and a high proportion (87.3%) of the genes were predicted to be pathogenic. Some diseases are highly related to copy number variations (CNVs), but there may be several genes involved when a responsible CNV is identified. We find the GPP score can significantly distinguish core genes from background genes of CNV, which may help us identify the core genes so that we can obtain accurate targets for later research.

There are some deficiencies in our study. When selecting non-pathogenic genes of MD (negative gene sets), we used the existing variants in public databases to select LOF variants tolerant genes. We didn’t find “reported non-pathogenic genes” because it’s hard to deem a gene to be non-pathogenic. This may produce some errors, although our analysis verified that the selected genes were generally accurate. When we applied the GPP score to analyze gene sets of different diseases, we selected only a limited number of diseases and genes. If more diseases and genes had been involved, we might have obtained more convincing results. We also tried to perform the GOF and LOF prediction by the same method, but we did not collect enough GOF genes. We may finish this effort to complement our study when we obtain more GOF genes. Furthermore, we think there is heterogeneity for different kinds of disease, so distinguishing the pathogenic genes for specific diseases, for example, ophthalmic diseases, neurological disease and developmental diseases, may be a new research direction.

In conclusion, our study estimates the total number of MD genes. And we introduce the gene pathogenicity prediction (GPP) score, which may provide robust and reliable guidance for the identification of pathogenic genes in MD research. We also provide two additional gene-level scores that may suggest the dominant or recessive inheritance model of genes. In addition, our results may promote the understanding of MD genes.

---

**Supplementary Fig 1.**
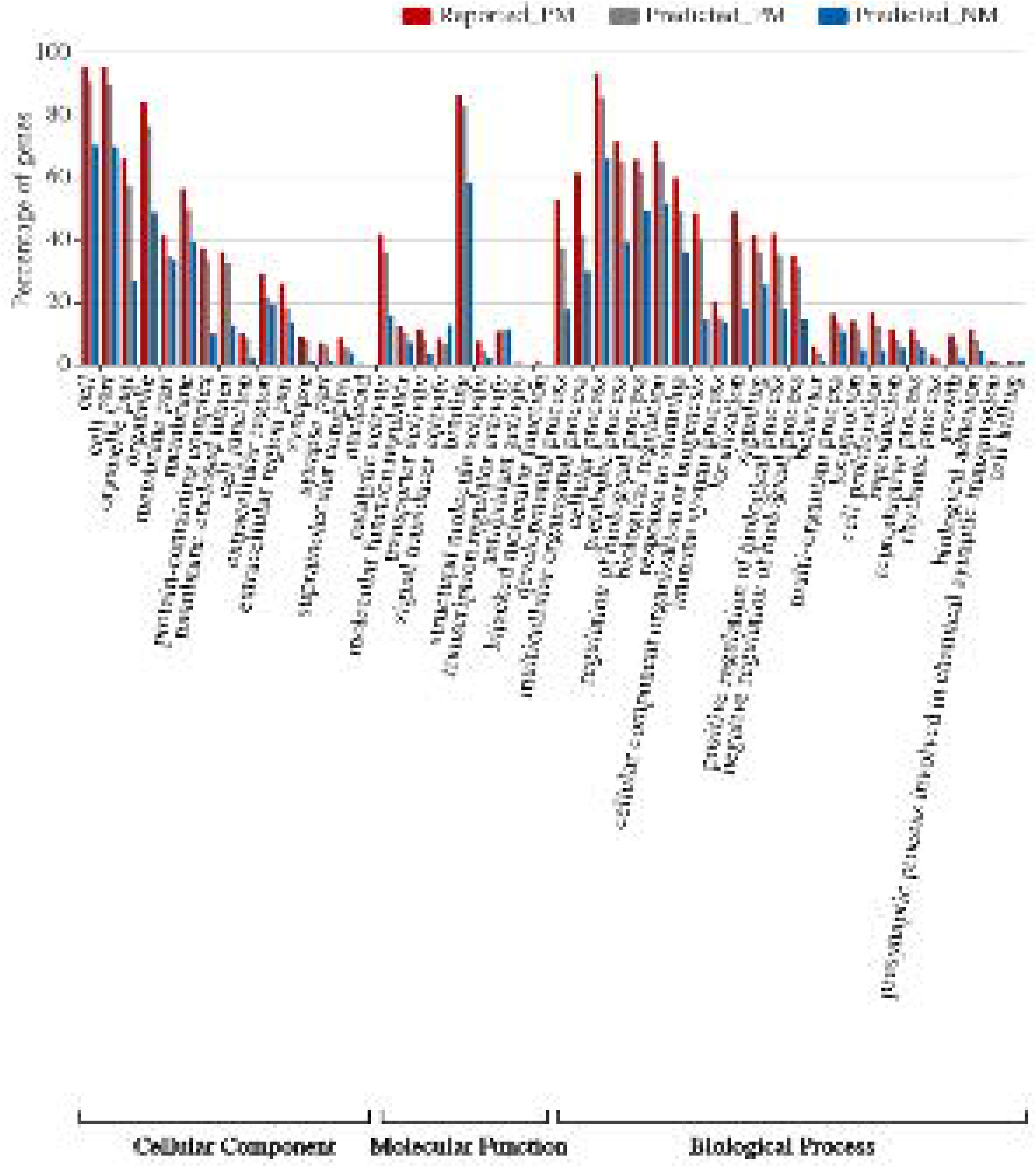
GO analysis of the predicted pathogenic and non-pathogenic genes of Mendelian diseases. The GO pathways analysis for reported_PM, predicted_PM and predicted_NM sets. The X-axis represents different GO pathways, while the Y-axis represents the percentage of genes in the selected pathway. The difference between the three gene sets are compared for each pathway. In general, the reported_PM and the predicted_PM sets show little difference, while the predicted_NM set show larger difference compared to them. Also, some opposite results are observed.

**Supplementary Fig 2.**
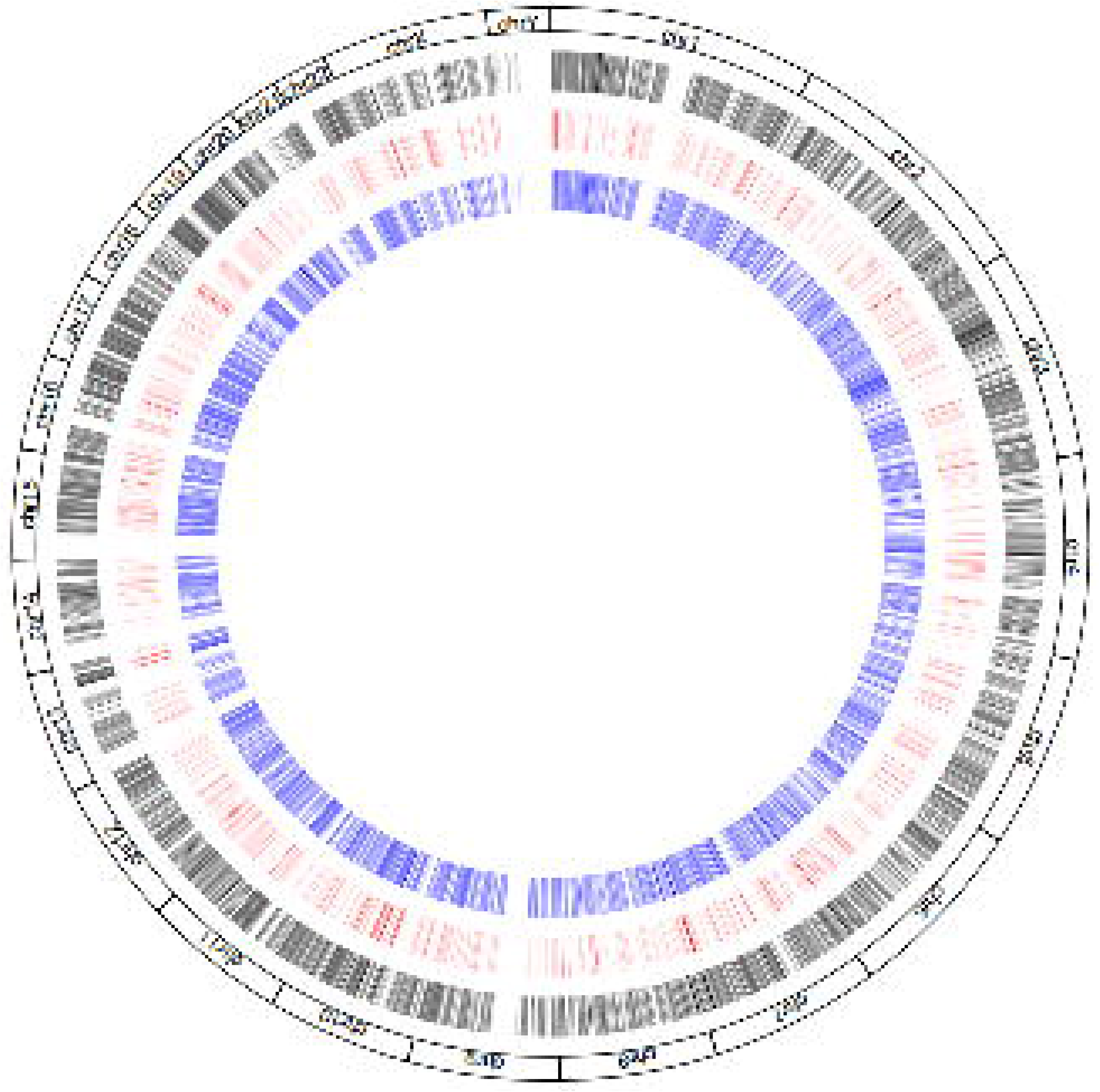
Distribution of predicted and reported Mendelian disease genes on each chromosome. The outer-layer circle colored black indicates the total protein-coding genes on each chromosome. The middle-layer circle colored red indicates the reported MD genes. The inner-layer circle colored blue indicates the predicted MD genes.

**Supplementary Fig 3.**
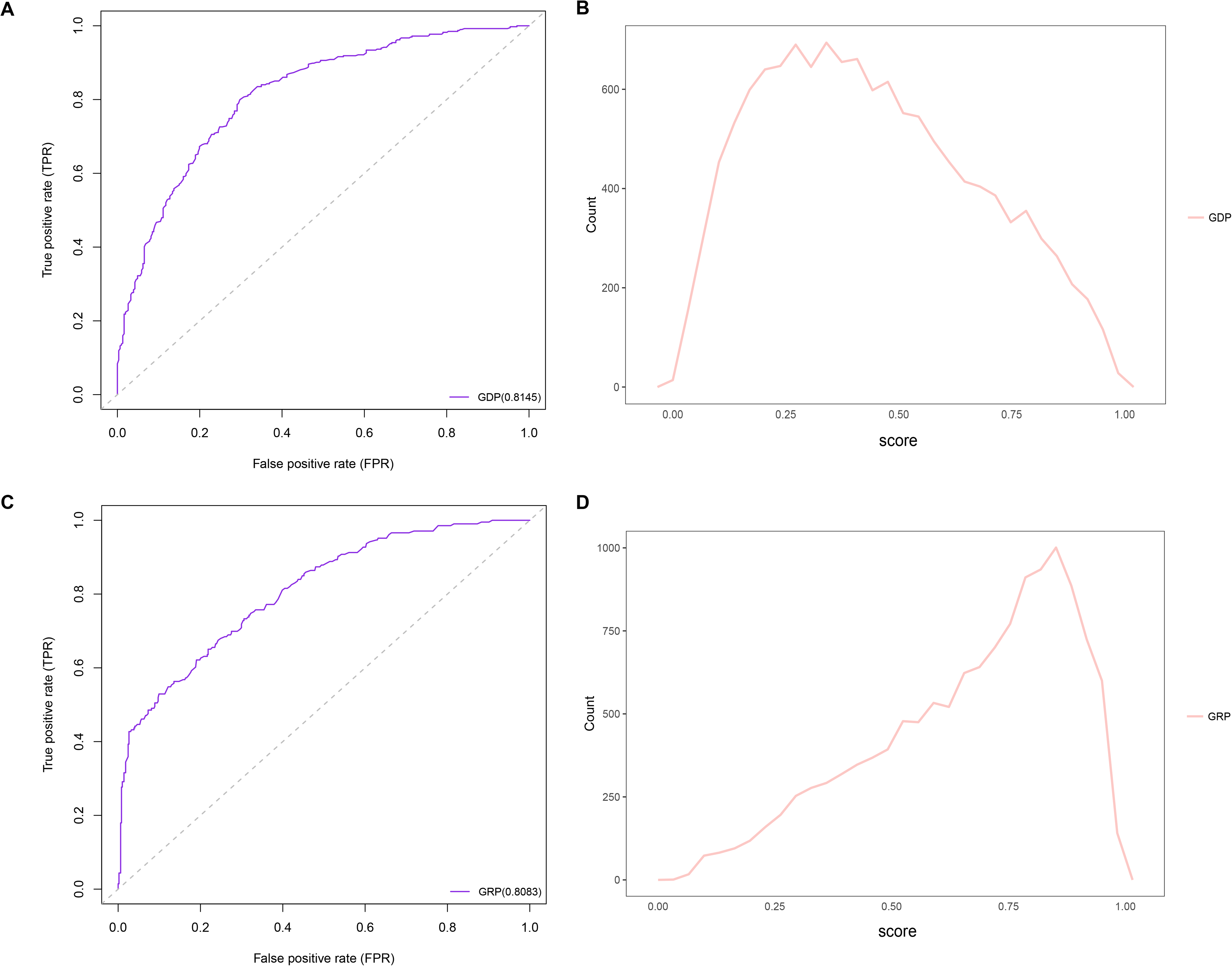
ROC curve of the GDP and GRP scores and the distribution of the scores of Mendelian disease genes. A. ROC curve of the GDP score. The AUC is 0.81. B. The distribution of the GDP score of MD genes indicates that fewer genes are dominant (the cutoff is 0.5). C. ROC curve of the GRP score. The AUC is 0.81. D. The distribution of the GRP score of MD genes indicates that more genes are recessive (the cutoff is 0.5).

## Acknowledgments

We thank the providers and maintainers of the public databases we used in our study. We appreciate the support of Shenzhen Key Laboratory of Neurogenomics (CXB201108250094A). This study was supported by the National Natural Science Foundation of China (81771444) and the Shenzhen Municipal Government of China (NO.JCYJ20170412153248372).

