## Supplementary material for "Gene pathogenicity prediction of Mendelian diseases via the Random Forest algorithm": table 1

**Table 1. Prediction accuracy of GPP score for several kinds of pathogenic gene set**

| **Gene set** | **Gene number** | **Percent of genes in training set (%)** | **Prediction accuracy (%)** |
| --- | --- | --- | --- |
| LOF intolerance genes | 3183 | 4.5 | 96.7 |
| Mendelian disease genes | 2946 | 20.4 | 92.8 |
| Pathogenic genes | 4893 | 12.7 | 87.4 |
| Susceptible genes | 98 | 0 | 74.5 |
