## supplementary method for "Gene pathogenicity prediction of Mendelian diseases via the Random Forest algorithm"

**Supplementary Methods**

**Data collection and gene standardization**

We obtained 19179 protein coding genes from HGNC (https://www.genenames.org/cgi-bin/statistics). As data from different sources may have inconsistent gene names, all the gene names were standardized in our study. If the gene name was consistent with the current HGNC symbol, it was retained. If the gene name was a previously used one, it was changed to the current HGNC symbol.

**Gene set selection and gene-level characteristic filtration in machine learning approach**

1) Gene set selection: Gene sets were collected through several approaches. a) We searched all pathogenic variants from ClinVar to find pathogenic genes. b) Disease-causing genes were obtained from OMIM, which included dominant genes, recessive genes, and susceptibility genes. c) Some genes for early-onset severe diseases were obtained from the “super hero project”. d) Some disease-causing genes were obtained from DOMINO, which included dominant genes and recessive genes. e) LOF variant tolerant genes were obtained from the KG, GAD and ExAC databases. We combined the genes obtained in parts a-d to build the reported pathogenic gene pool (5032 genes), and the genes from e were used as non-pathogenic genes (1480 genes).

We used the non-pathogenic genes shared among the 3 databases as the training set of negative genes (630 genes) because we considered them to have high quality, and the remaining ones were used as the testing set of negative genes (850 genes). To ensure balance in gene number between positive genes and negative genes used in our model, we extracted genes with clear inheritance models from OMIM (2984 genes) and randomly selected 1480 of them to be the positive gene set. In this process, we scored all the OMIM genes by RVIS, randomly selected them 3 times, and compared the difference between the selected genes and the rest genes by the Wilcoxon rank sum test. We then chose the ones with the highest p-value as the final ML pathogenic gene set (p=0.3328). Then, we used the same method to produce the training set of positive genes (630 genes) and the testing set of positive genes (850 genes) from the final positive gene set (p=0.7). We performed 5X cross-validation to ensure the uniformity of the gene sets.

We also tried two other positive gene set selection trials. In one, we used all 2984 genes and randomly selected half of them as the training set and the other half as the testing set. In the other, we used genes of complete penetrance, early-onset and severe Mendelian diseases as the training set (860 genes) and selected some high-quality genes of Mendelian diseases as the testing set (772 genes). The gene pathogenicity prediction results of the three approaches were quite similar, so we ultimately chose to use the first gene set selection strategy.

2) Gene level characteristic filtration: Some gene-level characteristics were selected and modified: a) We applied ANNOVAR to annotate variants from KG, GAD and ExAC with the dbnsfp30a package, which provided a number of variant damage prediction results from different software. b) We calculated the UTR length and CDS length for each gene with the gff file from Refgene. c) Gene-gene interaction data were obtained from STRING. d) Data on the conservation of nucleotides and amino acids between human and other species were obtained from HomoloGene. e) The expression data of different tissues were obtained from GTEx. f) The GC contents of the exons of each gene were calculated. g) Several gene-level characteristics selected by DOMINO were also included.

We also calculated the gene potential damage (GPD) score and gene intolerance scores by ourselves. We simulated all possible variants in exons of each gene and scored them by CADD. We then calculated the number of damage variants and the average, medium, maximum and minimum value of the GPD score of each gene. The calculation method of gene intolerance score was as follows:

Variants from the KG, GAD and ExAC databases were extracted and annotated. They were divided into common variants and rare variants based on the frequency in the population (we compared several cutoffs and finally identified 0.005 as the most suitable value). Additionally, the variants were divided into synonymous mutations (syn), missense mutations (mis), nonsense mutations (non), frameshift mutations (fs) and splice site mutations (splice). The syn category was regarded as “neutral mutations”, while the mis, non, fs and splice categories were regarded as “damaging mutations” even though some of them may not cause any negative effects. We calculated several scores for different variant sets separately.

We assumed that common variants had undergone natural selection, so the reserved variants should have no strong negative effects. Thus, the genes that harbored more “damaging mutations” tended to have functional tolerance. Because different genes may carry different numbers of variants, we used synonymous mutations for standardization when calculating the intolerance score. We calculated the number of mutations harbored by each gene separately for different kinds of mutation. For each kind of “damaging mutation”, the number was represented by Nh, the number for syn was represented by Ns, and the gene intolerance score was represented by S1: S1=Nh/ (Nh+Ns).

For rare variants, the situation seemed to be somewhat complicated, so we calculated the intolerance scores by 3 different methods.

a) We assumed some of the rare variants had undergone natural selection like common variants, so we applied the same method as for common variants, thus producing S2.
b) We also assumed some of the rare variants had occurred recently, so we used the mutation probability per gene per generation for each kind of mutation to calculate the expected number of mutations and compared it to the observed number of mutations. The mutation probabilities for different kinds of variant were obtained from DNE. The genes under strong selection tended to retain fewer “damaging mutations” than expected.

For each gene, we took the mutation probability of syn as x and the number of rare syn in the database as y. Then, we constructed a linear regression model for x and y. For each kind of “damaging mutation”, we applied this model with the mutation probability as x and then calculated the expected number of “damaging mutation”. We then compared the expected number (Ne) with the observed number (No) in the database to evaluate the intolerance score S3: S3=No/ (No+Ne).

c) Influenced by environment, when the variants passed from generation to generation, the “damaging mutation” likely transformed from common to rare because of selection. Therefore, the genes harbored more rare variants compared to common variants tend to be functional intolerance.

For each gene, we took the number of rare variants as Nr and the common variants as Nc. The transformation score (S) was calculated as S=Nr/ (Nr+Nc). Then, we used syn for standardization as above, which produced S4 for each kind of “damaging mutation”.

The 4 scores represented different gene-level characteristics. S1 reflected the density of common variants, S2 reflected the density of rare variants, S3 reflected the difference between observed and expected variants, and S4 reflected the difference between common and rare variants. A score was eliminated from the subsequent analysis if it contained less than 50% of all protein-coding genes. In total, we produced 48 gene intolerance scores (4 score calculation methods, 3 variant databases and 4 “damaging mutation” categories).

We included all the gene-level characteristics mentioned above. In total, 405 characteristics were enrolled, and we extracted all characteristics for each protein-coding gene. We excluded genes with missing values for more than 50% of all characteristics and we used the median value of other genes to fill in the remaining missing values. Then, we excluded characteristics that failed to distinguish positive genes from negative genes of the training set (ANOVA, p>0.05). If two characteristics were highly relevant (correlation coefficient >0.9), the better one was retained. Finally, 201 characteristics were left.

**Gene pathogenicity prediction score produced by random forest algorithm**

When gene sets and gene-level characteristics were obtained, we adjusted the parameters of the random forest algorithm. We tried 2 ntrees (ntree=500 and ntree=800) and several mtrys to select the best parameter values. We found that the two ntrees performed highly similarly, so we chose the default set (ntree=500). We added the accuracy and recall of the model (both for training set and testing set) and selected the top 5 mtry. Then we calculated the AUC of each mtry on the testing set and chose the best one as the final mtry.

After the parameter adjustment, we applied the machine learning approach to the training set and testing set. The importance of the characteristics was evaluated. We obtained the 10 most important characteristics for the model by the mean decrease in the Gini coefficient. Then, we predicted the pathogenicity of all protein-coding genes. After that, we collected different kinds of pathogenic gene sets, by which we further evaluated the accuracy of our GPP results. We further explored the distribution of the GPP score and the proportion of reported Mendelian disease genes for different scores, as well as the relationships between the GPP score and the severity and age of onset of diseases.

**Prediction of dominant and recessive models of Mendelian disease genes**

For the prediction of dominant and recessive genes, we extracted genes from OMIM and divided them into dominant, recessive, and both inheritance models. The genes on the X and Y chromosomes and mitochondrial were excluded. When we calculated the GDP score, the dominant and both inheritance sets were used as cases (1243 genes), while the recessive set was used as controls (1584 genes). When we calculated the GRP score, the recessive and both inheritance sets were used as cases (1985 genes), while the dominant set was used as controls (842 genes). Total characteristics were those used in the GPP score calculation. The filter strategy was the same. Finally, 183 characteristics were retained for the GDP score and GRP score calculations. We did not divide the genes into training and testing sets, and we applied 4X cross-testing for the calculation. Briefly, we randomly divided positive (case) and negative (control) sets into 4 equal groups. For each trial, we used 3 groups as the training set and 1 group as the testing set and repeated this process 4 times to ensure that all genes would be used as training and testing sets equally. Then, we applied the best-performing trial to build the model and calculate the scores.
